# Asymmetric chromosome segregation and cell division in DNA damage-induced bacterial filaments

**DOI:** 10.1101/2020.03.16.993485

**Authors:** Suchitha Raghunathan, Afroze Chimthanawala, Sandeep Krishna, Anthony G. Vecchiarelli, Anjana Badrinarayanan

## Abstract

Faithful propagation of life requires coordination of DNA replication and segregation with cell growth and division. In bacteria, this results in cell size homeostasis and periodicity in replication and division. The situation is perturbed under stress such as DNA damage, which induces filamentation as cell cycle progression is blocked to allow for repair. Mechanisms that release this morphological state for re-entry into wild type growth are unclear. Here we show that damage recovery is mediated via asymmetric division of *Escherichia coli* filaments, producing short daughter cells with wild type size and growth dynamics. Division restoration at this polar site is governed by coordinated action of divisome positioning by the Min system and modulation of division licensing by the *terminus* region of the chromosome, with MatP playing a central role in this process. Collectively, our study highlights a key role for concurrency between chromosome (and specifically *terminus)* segregation and cell division in daughter cell size maintenance during filamentous divisions and suggests a central function for asymmetric division in mediating cellular recovery from a stressed state.

## Introduction

For successful cell division to occur, accurate DNA duplication and segregation must be completed. In bacteria, chromosome replication initiates bi-directionally from an ‘*origin’* and finishes opposite to this position, at the ‘*terminus*’ (Reyes-Lamothe and Sherratt, 2019; Kleckner et al., 2014, 2018). Several factors ensure that cells divide only upon completion of this process by regulating the multiprotein division machinery called the ‘divisome’ (Galli and Gerdes, 2012; Dewachter et al., 2018; Kleckner et al., 2018; Männik et al., 2016). For example, *E. coli* encodes negative regulators of the tubulin homolog FtsZ, that is required to initiate the assembly of the divisome at the division plane. Nucleoid occlusion by SlmA prevents the formation of the FtsZ-ring at locations where chromosomal DNA is present and MinCDE oscillations direct the position of the Z-ring near mid-cell (Tonthat et al., 2011; Bernhardt and de Boer, 2005; Tsang and Bernhardt, 2015). Recent studies have also suggested coordination of division with the *terminus* via proteins such as MatP and ZapAB that act as a bridge between the DNA as well as the divisome (Espéli et al., 2012; Mercier et al., 2008; Männik et al., 2016). Together, in unperturbed laboratory conditions, this results in daughter cells that replicate and divide in a periodic manner and that do not show much deviation in birth and division cell sizes (Donachie, 1968; Taheri-Araghi et al., 2015; Wallden et al., 2016; Micali et al., 2018; Campos et al., 2014; Si et al., 2019; Harris and Theriot, 2016). Such size maintenance has been described in other bacteria as well as eukaryotic systems (Chandler-Brown et al., 2017; Lambert et al., 2018; Soifer et al., 2016).

Recent investigations in bacteria have proposed that cells follow an ‘adder’-based principle that governs size homeostasis either via regulation at the stage of birth (replication initiation) or division or both (Taheri-Araghi et al., 2015; Wallden et al., 2016; Campos et al., 2014; Si et al., 2019). In addition, some studies have also proposed a role for cell shape or concurrency between processes of replication and division in size control (Micali et al., 2018; Harris and Theriot, 2018). This homeostatic state is perturbed under conditions of stress, a situation that can be often faced by bacterial cells in their environment (Horvath et al., 2011; Justice et al., 2008; Yang et al., 2016; Heinrich et al., 2019). For example, under DNA damage, a cell cycle checkpoint blocks cell division until damage has been repaired. The bacterial SOS response is activated upon the binding of RecA to single-stranded DNA that is exposed as a consequence of DNA damage (Mukherjee et al., 1998; Jonas, 2014; Kreuzer, 2013). As part of this response, a cell division inhibitor (such as SulA in *E. coli*) blocks polymerization of FtsZ, resulting in cellular elongation or filamentation (Kreuzer, 2013; Mukherjee et al., 1998). Cell division inhibition during damage is a conserved process, even though the effectors may vary across bacteria. SOS-induced division inhibition is carried out by SidA in *Caulobacter* and YneA in *Bacillus* (Mukherjee et al., 1998; Jonas, 2014; Modell et al., 2011; Mo and Burkholder, 2010). SOS-independent DNA damage-induced division inhibitors have also been identified, suggesting that this is an important step in DNA repair (Modell et al., 2014). Along with blocking division, chromosome cohesion or de-condensation is also initiated in these filaments. It is thought that this can aid recombination-based repair (Vickridge et al., 2017; Odsbu and Skarstad, 2014). Indeed, cells have also been shown to change shape and size under other forms of stress including host environments, heat shock and osmotic fluctuations (Justice et al., 2008; Heinrich et al., 2019; Wehrens et al., 2018; Kysela et al., 2016; Caccamo and Brun, 2018; Bos et al., 2015; Yang et al., 2016). Together, this highlights the plasticity with which bacteria such as *E. coli* sample a range of cell sizes including filamentous and non-filamentous cell lengths. While the process by which cell division is regulated to result in elongation under DNA damage has been well-characterized (Mukherjee et al., 1998; Suzuki et al., 1967; Jonas, 2014; Modell et al., 2014, 2011; Mo and Burkholder, 2010; Kantor and Deering, 1966; Adler and Hardigree, 1965), how such a state is exited after repair to reinitiate wild type, periodic replication and division remains unclear.

In this study, we probe the mechanism by which filamentous *E. coli* reinitiate chromosome segregation and cell division after DNA repair. We use single-cell, time-resolved fluorescence microscopy to follow the kinetics of division restoration after cells face a pulse of DNA damage and observe that size is maintained in daughter cells generated from dividing filaments. Size homeostasis in daughters is not governed by the growth laws (in the filament) and is instead dictated by cell division. We further find that division restoration is controlled by two steps: determining *where* and *when* to divide. This stepwise process, regulated by a combination of MinCDE oscillations, FtsZ levels and *terminus* segregation, is accompanied by asymmetric partitioning of repaired chromosomes, resulting in the production of daughter cells of the right size and devoid of DNA damage, thus facilitating recovery from a stressed state.

## Results

### DNA-damage induced filaments maintain daughter cell size during division

To understand how damage-induced filamentous *E. coli* (Supplementary video 1) re-initiate cell division and wild-type growth after DNA damage, we followed division restoration in cells after treatment with a sub-inhibitory dose of the DNA damaging agent, Mitomycin-C via time-lapse imaging (1μg/ ml; (Dapa et al., 2017)) (Fig. 1A). While unperturbed wild type cells divided at ∼mid-cell (Fig. 1B), we found that a significant proportion of cells deviated from this division pattern as they increased in length (Fig. 1C-D). Damage-treated cells close to wild type length (5-10 μm) tended to divide in the middle resulting in production of two daughter cells of similar sizes. In contrast, filamentous cells divided polarly (or asymmetrically) to produce a short ‘daughter’ cell (S_D_) of size similar to wild-type and a long cell that continued to filament (L_D_) (Fig. 1C and 1E; Supplementary video 2). The probability of a cell to undergo asymmetric division increased with increasing cell length with 85% cells dividing asymmetrically at lengths >12μm. Varying durations of damage exposure (30, 60 or 90 min) resulted in different degrees of filamentation. However, in each case, a filamentous cell at division resulted in production of an S_D_ that was close to wild type size (Fig. 1F-G and Fig. S1A-B). This asymmetry in division was observed in damage-induced filaments independent of growth media (Fig. S1C). To further test whether the switch from mid-cell to asymmetric division was a consequence of filamentation, we treated *E. coli* with the cell division inhibitor, cephalexin, where genome integrity is maintained and chromosome replication and segregation takes place as wild type. However, cells are unable to divide, resulting in elongation in length (Rolinson, 1980; Chung et al., 2009). Cephalexin-treated filamentous cells also generated daughter cells (S_D_) of fixed length via asymmetric division upon removal of the inhibitor (Fig. S1D; Supplementary video 3). Together with previous observations (Wehrens et al., 2018; Taschner et al., 1988; Begg and Doanachie, 1977), our data suggest that filamentous cells, irrespective of types or durations of stress treatment, engage in this form of division that produces daughter cells of wild type size.

**Figure 1:**
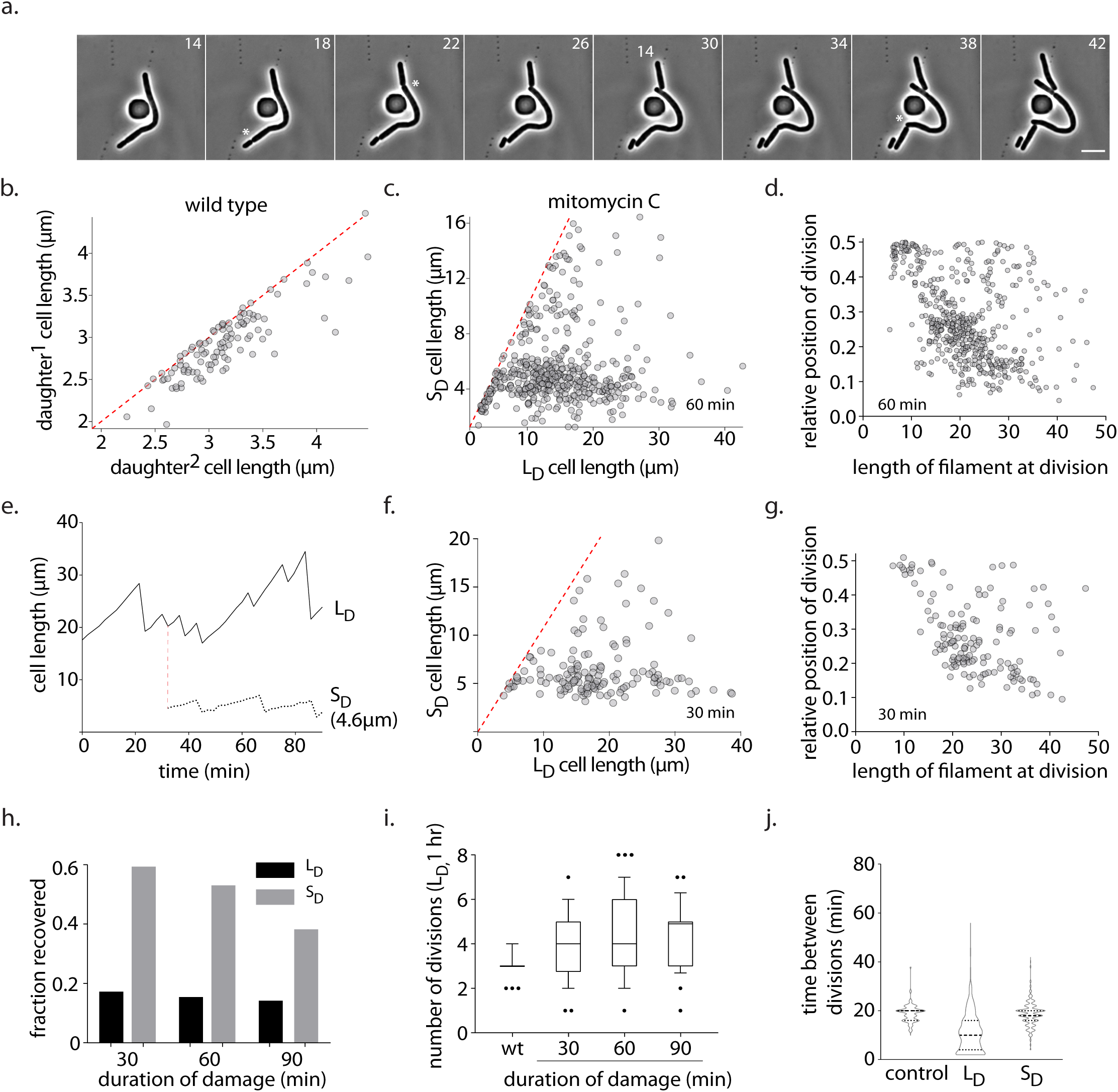
DNA-damage induced filaments maintain daughter cell size during division. a. Representative time-lapse montage of filamentous cells during recovery. White asterisks indicate divisions occurring towards a cell pole. Scale bar - 5 µm; time in min here and in all other images. b. Cell length of two daughter cells generated from a single division in wild type conditions. Each grey dot represents a single division event (n = 157). The red line plots the expected values if all cells were dividing at their mid-point. c. Cell length of long daughter (L_D_) and short daughter (S_D_) generated from a DNA damage-induced filament during recovery. Cells are treated with mitomycin C (MMC) for 60 min. Each grey dot represents a single division event (n = 531). The red line plots the expected values if all cells were dividing at their mid-point. d. Location of division is plotted as a function of cell length in filamentous *E. coli* during recovery from DNA damage treatment (60 min; n = 531). e. Cell length of long daughter (L_D_) and short daughter (S_D_) is tracked over time during damage recovery. Decrease in cell length is indicative of division. f-g. As (b-c) for cells treated with MMC for 30 min. h. Fate of S_D_ and L_D_ during recovery. Cell is classified as recovered if it undergoes mid-cell division and filamentous if it continues to filament after division (n ≥ 56) i. Number divisions per cell in 1 hr for all durations of damage treatment. As a control, number of divisions wild type cells undergo is also shown (n≥100 division). j. Distribution of time between divisions for wild type (no damage control), L_D_ and S_D_ during recovery from MMC (n ≥ 104).

To characterize the recovery process further in DNA damage-induced filaments, we followed the fate of the filament (L_D_) and the S_D_ over time. We observed that only 16±2% of filaments returned to wild type cell size during recovery in the three treatment regimens (Fig. 1H). However, filamentous cells underwent multiple divisions in a 1 hr time period, generating daughter cells (S_D_) of wild type size at each division (Fig. 1I). In the same time, wild type cells undergo three divisions on average, suggesting that more daughter cells are produced from a filament than a cell of same size as wild type during recovery. In contrast to the filaments, daughter cells of wild type size (S_D_) displayed growth and division dynamics similar to non-damage conditions (Fig. 1H and 1J). As an example, in Fig. 1E, we followed the fate of an S_D_ (4.6 μm) generated from a filament and found that the daughter cell reinitiated wild type growth dynamics soon after division. Time taken between divisions for S_D_ was close to the distribution seen for wild type, while L_D_ continued to grow and divide as damage-treated cells (Fig. 1J). Consistently, the DNA damage marker, RecA, formed multiple foci in L_D_ at the time of division, while S_D_ had RecA localization as seen for wild type cells (Lesterlin et al., 2014; Rajendram et al., 2015; Vickridge et al., 2017) or elongated cells with no DNA damage (cephalexin treated) (Fig. S1E). In line with these observations, we found that viable cell count as well as cell length distribution was restored to close to that of wild type in the three treatment regimes (Fig. S1F-G). Thus, even though the filament may or may not recover from damage exposure, recovery of the population may be mediated via several rounds of asymmetric divisions from a single filament, resulting in daughter cells of wild type size and growth dynamics.

### Daughter cell size is governed by cell division dynamics of filaments

How is daughter cell size maintained in dividing filaments? Recent investigations have extensively characterized various mechanisms for chromosome segregation checkpoints and adder-based cell size maintenance principles in wild type cells (Taheri-Araghi et al., 2015; Campos et al., 2014; Si et al., 2019; Harris and Theriot, 2016; Arjes et al., 2014; Kleckner et al., 2014; Hill et al., 2012; Campos et al., 2018). To address this question in the current context, we analysed the growth and division dynamics of filaments. In damage-induced filaments, we found that length added between divisions was not fixed (Fig. 2A) and did not correlate with length of the cell at birth, with several instances of consecutive divisions without any elongation in between each division. To test whether there is periodicity in division timing, we measured time between consecutive division events. While wild type cells divided every ∼20min, filamentous cells displayed a large distribution of times between consecutive division events (Fig. S2A). Although a lack of periodicity in division timing was observed, we found that 67±4% of divisions took place at the opposite pole of the previous division, suggesting that there may be no preference for old or new pole during division licensing (See Fig. S2D-G and materials and methods for details on how divisions are computationally analysed).

**Figure 2:**
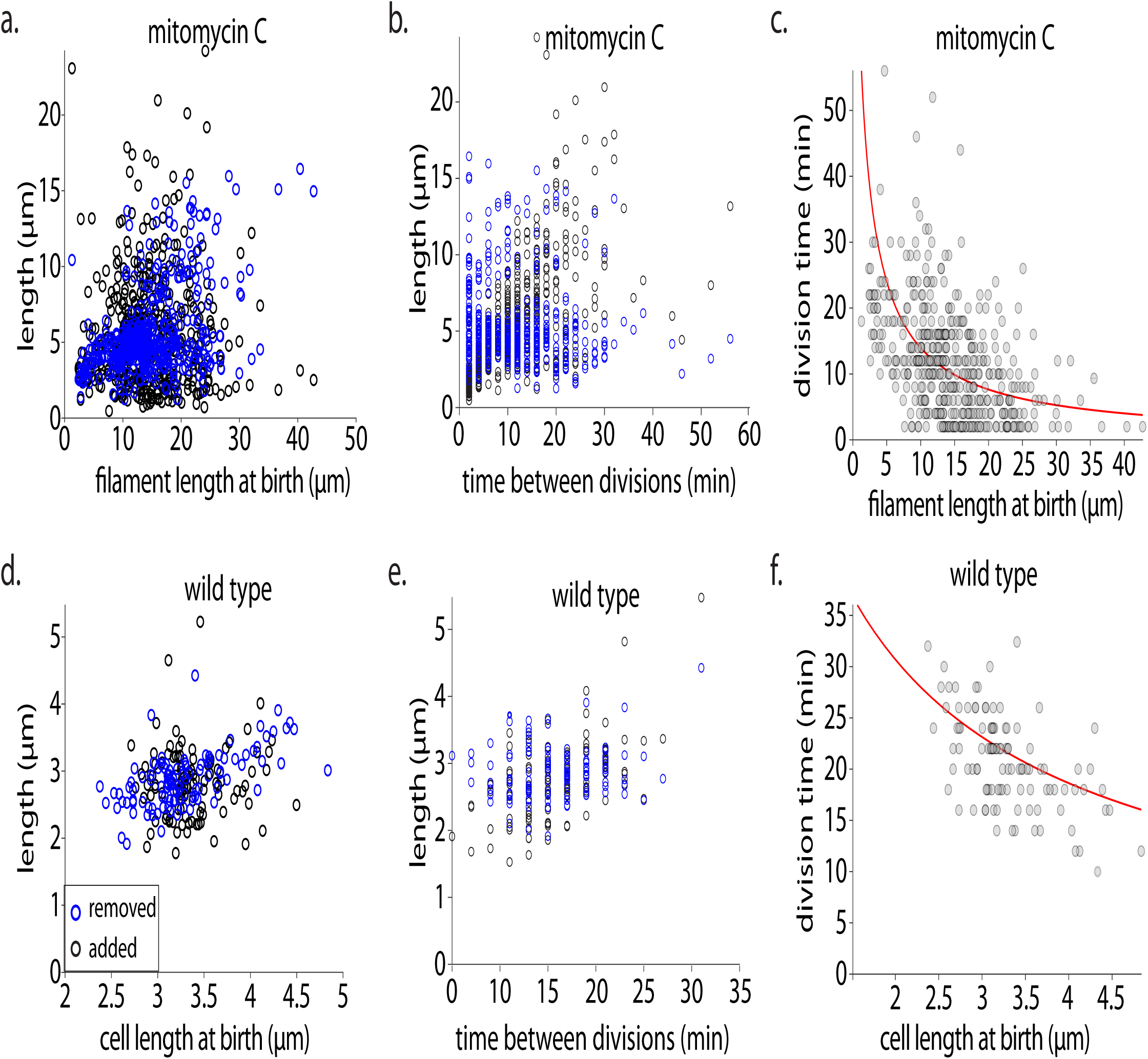
Daughter cell size is governed by cell division dynamics of filaments. a. Length added to a cell between divisions (black dots) and length removed at the latter division (i.e., the length of the daughter cell) (blue dots) as a function of the birth length of the cell for MMC-treated cells. b. Length added to a cell between divisions (black dots) and length removed at the division (i.e., the length of the daughter cell) (blue dots) as a function of the time between divisions for MMC-treated cells. c. Time between divisions as a function of cell length at birth. Red line shows the relation that would be necessary for the system to be an adder (equation (3) in supplementary results), given that cells are growing exponentially with the rates given in Fig. S2B-C. d-f. (As a-c) for wild type cells.

After damage filament length increased exponentially with a characteristic rate, as seen in wild type conditions (Fig. S2B-C), with some fluctuation which probably arises because of stochasticity in damage and/ or repair. When length added or removed was plotted as a function of birth length in filaments, no clear trend was observed (Fig. 2A). However, when length added/ removed was plotted as a function of division times, we observed that length removed (S_D_ cell length) neither increased nor decreased with inter-division time, while length added increased as time between divisions increased (Fig. 2B). This is in contrast to wild type cells where length added/ removed are constant (Fig. 2D-E). Overall, our observations suggest that the adder mechanism seen in steady state conditions breaks down in filamentous cells undergoing asymmetric division (see methods section for analysis). This could occur if growth rate changes, or if the relationship between time to division and birth length deviates from what is expected (eq. 3 in supplementary results for a model of this relationship), or both. In case of elongated cells, we find that cells continue to grow exponentially at a characteristic rate regardless of length, while the relation between time to division and birth length becomes much more noisy. Indeed, when we plotted division time as a function of birth length, a decreasing trend was most clearly observed only for wild type and less so (with more noise) in the case of damage-treated cells (Fig. 2C and 2F). However, two properties of filament division remain constant: a. length removed (S_D_ cell size) is invariant and b. division takes place poleward, asymmetrically. Taken together, this supports the idea that daughter cell size is governed by regulation of cell division rather than filament growth dynamics.

### Cell division positioning and licensing have distinct regulatory mechanisms

#### Role of Min system in divisome positioning

As stated above, we noticed that cells transitioned from mid-cell to asymmetric divisions as they grew longer in length (Fig. 1C). To test if the Min oscillation system (Wehrens et al., 2018; de Boer et al., 1989; Raskin and de Boer, 1999; Tsang and Bernhardt, 2015; Bisicchia et al., 2013; Dewachter et al., 2018) could determine division licensing events in damage-induced filaments, we first asked whether Min oscillations are maintained in these long cells. We imaged MinD-GFP and, consistent with Wehrens et al., 2018, found that it localized at several positions along the length of the cell and in an equidistant manner, with a distance of 7 µm on average between each Min localization in MMC or cephalexin-treated cells (Fig. S3A). As expected, we observed that cells had 1-2 Min localizations at lengths between 5-10µm and the number of Min localizations increased with increasing cell length (Fig. S3B). Multiple Min localizations would suggest that the number of divisome localizations would also proportionally increase. However, when we imaged the localization of FtsZ (Supplementary video 4) in filaments, we found that the number of FtsZ rings did not scale with increasing cell length and the position of FtsZ shifted from away from mid-cell as cell length increased (Fig. S3C). Importantly, deletion of the *min* operon resulted in loss of cell size maintenance with divisions that could occur at mid-cell as well (Fig. 3A and S3E). Thus, as in wild type conditions, the Min system may dictate daughter cell size maintenance in damage-induced filaments as well.

**Figure 3:**
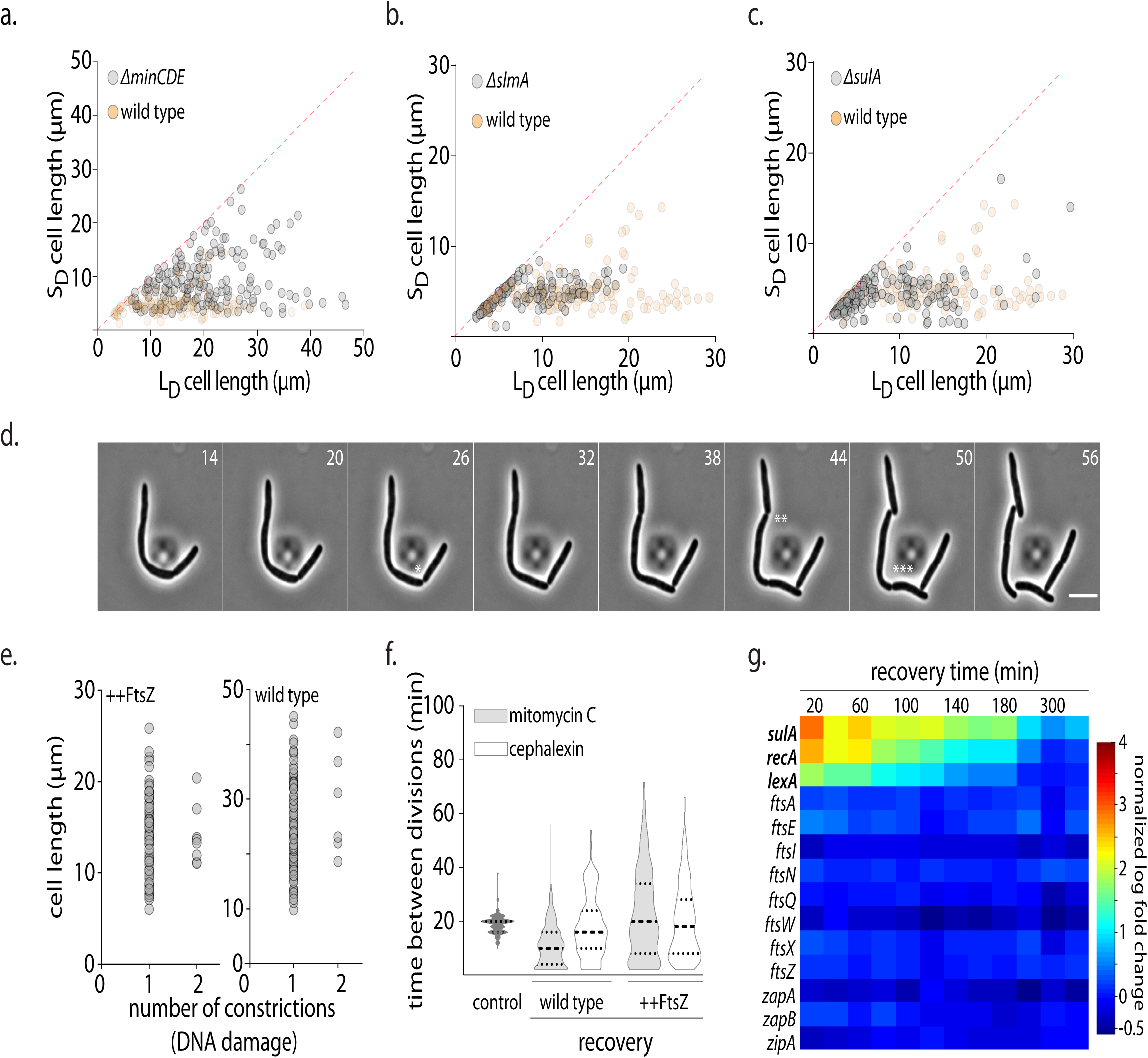
Cell division positioning and licensing have distinct regulatory mechanisms. *Determining where to divide: role of Min system.* a-c. Cell length of long daughter (L_D_) and short daughter (S_D_) generated from a DNA damage-induced filament during recovery for *ΔminCDE* (n=186), *ΔslmA* (n=144) and *ΔsulA* (n=246) backgrounds respectively (grey dots). As a reference, lengths for wild type during recovery are shown in orange. The red line plots the expected values if all cells were dividing at their mid-point. d. Representative time-lapse montage of division in wild type cells during damage recovery. e. Number of constrictions per cell plotted as a function of cell length during DNA damage recovery for wild type and cells over-expressing FtsZ (n ≥ 91). f. Time between divisions for no damage (control) and for cells after treatment with MMC or cephalexin with (++FtsZ) or without (wild type) FtsZ overexpression (n ≥ 93). g. Heat map of transcript levels (from RNA-seq) of genes involved in cell division during a damage recovery time-course. As a control, genes induced under the SOS response are also highlighted (bold). Log_2_-fold change normalized to control without damage is plotted.

Even though daughter cell size maintenance was inaccurate in a *min* deletion, we found that in both wild type and *Δmin* cells division occurred only one-site-at-a-time (Fig. 3D, S2D-G and S3E). Since we did not observe multiple FtsZ rings or multiple constrictions in recovering filaments, we wondered whether nucleoid occlusion or negative regulation of FtsZ polymerization (Kreuzer, 2013; Mukherjee et al., 1998; Tonthat et al., 2011; Bernhardt and de Boer, 2005) may play a role in restricting the numbers and hence timing of division, resulting in division occurring one-site-at-a-time. However, we found that daughter cell length is maintained in *sulA* or *slmA* deletions and the number of constrictions did not increase (Fig. 3B-C and S3D). We then wondered whether FtsZ levels itself could be limiting in these cells (Bi and Lutkenhaus, 1990), thus permitting only one division event at a time. To test this, we over-expressed FtsZ from an arabinose-inducible promoter during damage or cephalexin recovery. In the case of damage-induced filaments, we did not observe more than one constriction on average in both wild type and FtsZ overexpression conditions, irrespective of the length of the cell (Fig. 3E and S3F). Given that chromosome segregation is perturbed in these conditions, we reasoned that the effect of FtsZ overexpression would be better observed in cephalexin-treated cells, where 30% cells had more than one constriction site in wild type background (Fig. S3D). We found that overexpression of FtsZ in cephalexin recovery resulted in the formation of multiple constriction sites in long cells (Fig. S2G, S3D and S3F). However, division timing did not become faster here or in the damage-recovery case (Fig. 3F) and division still occurred one-site-at-a-time (Fig. S2D-G and S3F). It is possible that another divisome component, such as FtsN (Coltharp et al., 2016), may be limiting in filaments, thus resulting in single division events. While we cannot rule out regulation of the divisome at the level of protein activity, transcriptional profiling of cells during recovery from MMC treatment did not show down-regulation of key division components (Fig. 3G). Thus, although the Min system governs division site locations, some other factor(s) determine division site licensing/ timing.

#### Regulation of division licensing: impact of chromosome and terminus segregation

Given that chromosomes in DNA damage-induced filaments are no longer segregated, it is possible chromosome dynamics in filaments could contribute to control of division licensing. To ascertain the role of chromosome segregation in regulation of division timing, we followed chromosomes after DNA damage via imaging the nucleoid-associated protein, HupA (Youngren et al., 2014; Marceau et al., 2011), tagged with mCherry after MMC or cephalexin treatment (Fig. 4A; Supplementary video 5). Since filamentous cells carry several copies of their chromosomes, yet division licensing occurs towards a cell pole, we wondered whether there could be differences in chromosome segregation dynamics across the length of the filament. Estimation of location of least intensity of HupA fluorescence (as a proxy for nucleoid-free regions and thus chromosome segregation) revealed that such regions were not near mid-cell in a majority of damage or cephalexin-treated elongated cells (Fig. 4B). In contrast to wild type, where median relative position of nucleoid-free gap from a cell pole was 0.46 (∼mid-cell), the location of this gap was broadly distributed between the cell pole and mid-cell with median position of 0.33 and 0.34 from a cell pole for MMC and cephalexin-treated cells respectively. Indeed, the location of the FtsZ-ring also coincided with where the chromosomes were most segregated at the time of division in our damage-recovery conditions (Fig. S4A-B).

**Figure 4:**
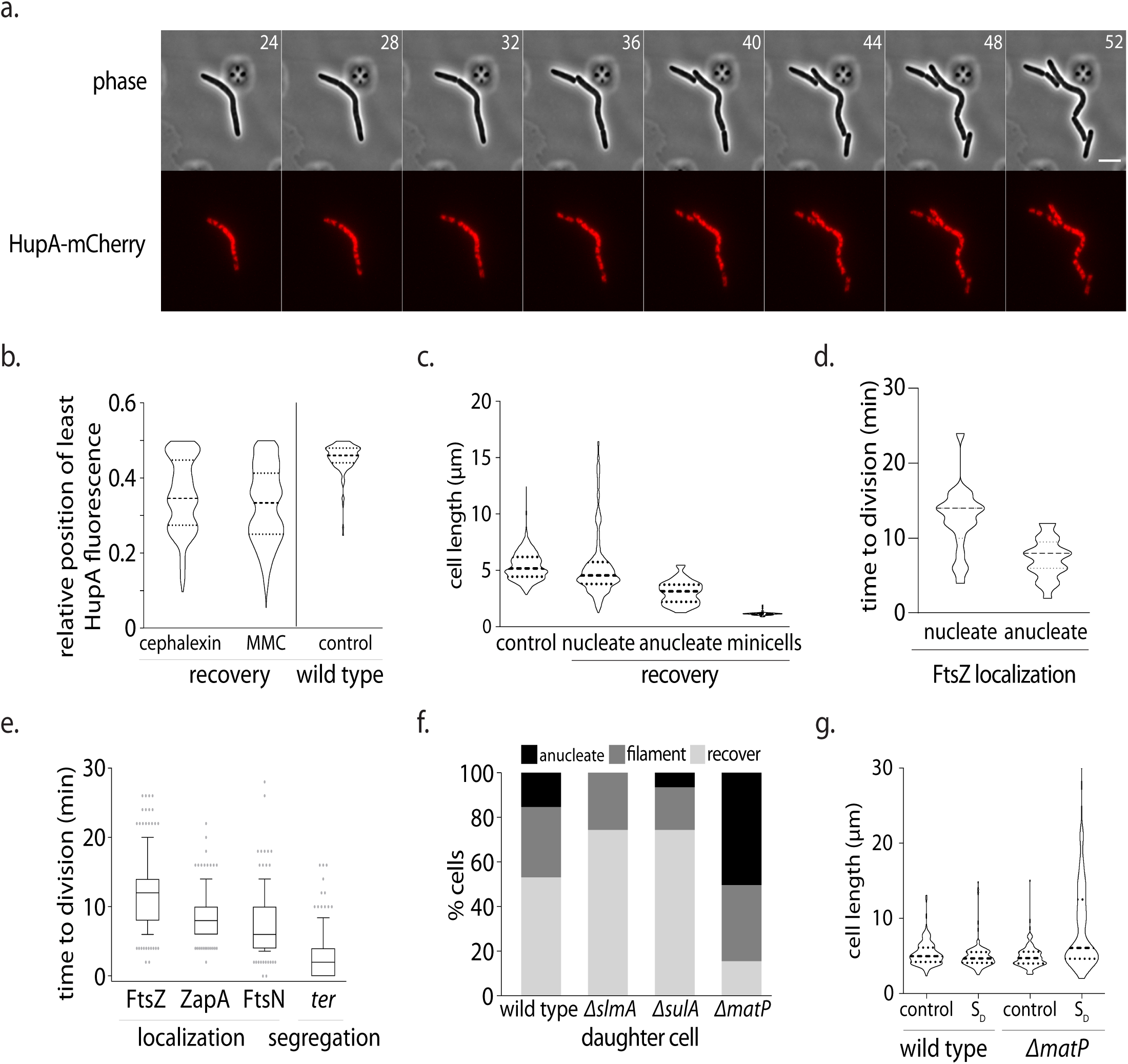
Cell division positioning and licensing have distinct regulatory mechanisms. *Regulation of division licensing: impact of chromosome and terminus segregation.* a. Representative time-lapse montage of division in cells during recovery. Grey – phase, red – HupA-mCherry (chromosome); scale bar - 5 µm; time in min. b. Position of least intensity of HupA fluorescence (gaps between chromosomes) plotted as a function of cell length (from one pole to mid-cell) in recovering MMC or cephalexin-treated filaments. As reference, these data are also shown for wild type cells with no damage treatment (control) (n ≥ 150). c. Cell length distribution for wild type cells (no perturbation) is plotted. Along with this, cell length distribution of daughter cells during DNA damage recovery is plotted for nucleate cells and anucleate cells. To highlight the distinction between these cell division events and minicell formation, cell length distribution of minicells (from *min* deleted cells) is also shown (n ≥ 100) d. Time from FtsZ localization to division completion is plotted for divisions that result in nucleated or anucleated S_D_ cells. (n ≥ 38) e. Distribution of time to division after FtsZ, ZapA and FtsN localization to division site is plotted. Along with this, time to division after segregation of *terminus* (*ter*) during recovery after MMC treatment is also shown (n ≥ 93). f. Percentage of S_D_ that are anucleate, recover and filament is plotted for wild type and deletions of *matP, sulA* or *slmA* during DNA damage recovery. (n ≥ 103) g. Daughter cell length distribution for nucleated divisions in the case of *matP* deletion and wild type is plotted. As a reference cell length distribution for cells without damage treatment is also plotted for each case (n = 145).

Interestingly, we observed that early divisions in long filaments resulted in 15±6% of daughter cells that were anucleated. These cells were distinct from mini-cells (Adler et al., 1967) in that their cell size was close to that of wild type (3.5µm) (Fig. 4C). We found that 38% of anucleate divisions occurred at the first division event during recovery. This percentage reduced to <7% by the third division. Thus, we hypothesized that if chromosome segregation has no effect on division dynamics then the time from divisome assembly to cell division should be the same between anucleated and nucleated divisions. However, if segregation had an effect, then the dynamics would vary between the two types of divisions. In the case of divisions that resulted in production of anucleated cells, we found that the average period between FtsZ assembly at the division site to cell division was 7 min. In contrast, in case of nucleated divisions, we noticed that FtsZ localized in the nucleoid-free regions for significantly longer, with an average of 12 min prior to division (Fig. 4D).

We wondered what could be the rate limiting step in allowing division to occur, thus resulting in persistence of FtsZ in nucleoid-free regions in the case of nucleated cell divisions. Given that the *terminus* region of the chromosome is the last to be segregated prior to division in wild type cells, we followed the dynamics of the *terminus* during division in recovering filaments using the *parS-*ParB locus labelling system (Espéli et al., 2012; Nielsen et al., 2006) (Fig. S4C-D). While the bulk of the chromosome segregated well-before division, and divisome components downstream of FtsZ, (ZapA and FtsN; Supplementary video 6-7), localized 8 min and 6 min prior to division respectively, a single *terminus* focus could be observed to persist in the chromosome-free region, where constriction had begun to occur. The constriction completed to division just prior to or concomitant with the *termini* splitting into two foci on either side of the division plane, within 2 min on average (Fig. 4E, S4C-D). If *terminus* segregation is indeed key to division licensing to generate nucleated, viable daughter cells, then perturbing this function should affect division dynamics during recovery. A key player in modulating *terminus* segregation specifically is MatP, which has been implicated in a. structuring the *ter* macrodomain (via prevention of MukBEF interaction with the terminus) and b. connecting the *ter* region with the divisome (Espéli et al., 2012; Lioy et al., 2018; Mercier et al., 2008; Nolivos et al., 2016). We found that a *matP* deletion was compromised in damage recovery (Fig. S4E). These cells had a significant increase in anucleate cell production, with 50±3% of divisions being anucleated. This is in contrast to wild type or deletions of *slmA* or *sulA* that do not affect *terminus* segregation, but instead regulate divisome assembly (Fig. 4F). In addition, nucleated daughter cell size maintenance was inaccurate in *matP* deleted cells (Fig. 4G). These data are consistent with the idea that chromosome and more specifically, *terminus* segregation acts as the licensor of nucleated divisions in filaments. Taken together, this suggests that coordination between Min-mediated divisome localization and MatP-mediated *terminus* positioning/ segregation regulates daughter cell size maintenance, thus facilitating recovery from DNA damage (Fig. S5).

## Discussion

In laboratory conditions, wild type *E. coli* maintains a distinct periodicity of cell growth and division that appears to be coupled with chromosome replication and segregation (Donachie, 1968; Taheri-Araghi et al., 2015; Wallden et al., 2016; Micali et al., 2018; Campos et al., 2014; Arjes et al., 2014; Kleckner et al., 2014; Hill et al., 2012). However, it is becoming increasingly evident that bacteria can exist in diverse morphological states, in part dictated by their environmental conditions (Yang et al., 2016; Justice et al., 2008; Heinrich et al., 2019; Jonas, 2014; Muraleedharan et al., 2018; Kysela et al., 2016; Caccamo and Brun, 2018; MacCready and Vecchiarelli, 2018). Even *E. coli* can become highly filamentous under conditions of stress such as during infection (Horvath et al., 2011; Justice et al., 2004). Transitions into filamentous morphologies are thought to confer several advantages such as avoiding phagocytosis via the host immune response or providing a means to dilute the effects of any inhibitors present in the surroundings (Yang et al., 2016; Justice et al., 2008; Horvath et al., 2011; Justice et al., 2004). Recent reports have also suggested that filamentation (at least in the case of *E. coli* treated with ciprofloxacin) may be the first step towards the emergence of antibiotic resistance as daughter cells carry mutations making them ciprofloxacin resistant (Bos et al., 2015). Thus, it becomes important to understand how bacterial cells enter and exit these filamentous states to ensure survival under stress. Indeed, several studies have characterized mechanisms by which filamentation is induced under DNA damage (Kreuzer, 2013; Mukherjee et al., 1998; Suzuki et al., 1967; Jonas, 2014; Modell et al., 2014, 2011; Adler and Hardigree, 1965; Uphoff, 2018). However, once DNA repair has been completed, how cells restore wild type growth is unclear.

Here we show that cell cycle restoration involves asymmetric chromosome segregation and cell division in filaments, resulting in the production of daughter cells of wild type cell size and with undamaged DNA. While the filament itself may or may not recover, several short daughters generated from a single filament go on to replicate and divide as wild type cells. The concept of asymmetric partitioning of cellular components during stress seems to have been co-opted by several bacterial systems (Schramm et al. 2020) including *M. tuberculosis,* where one daughter cell inherits the growing pole, while the other has to assemble a growth pole *de novo*. This results in difference in susceptibility to antibiotic treatment between the two cell types; the daughter cell with the growing pole is more sensitive to cell wall synthesis inhibitors when compared to cells that inherited the non-growing pole (Aakre and Laub, 2012; Aldridge et al., 2012). The swarmer cells of *P. mirabilis* and *V. parahaemolyticus* (Muraleedharan et al., 2018; MacCready and Vecchiarelli, 2018) use the Min-system to regulate asymmetric cell division to allow for such division while still preserving the population of filamentous cells. Dim-light stress induces filamentation in the photosynthetic cyanobacterium *S. elongatus*, which then divides asymmetrically via positioning by the Min system (Liao and Rust, 2018). Even in *E. coli* filaments generated in non-DNA damage conditions, studies have reported poleward division events (Adler and Hardigree, 1965; Wehrens et al., 2018; Taschner et al., 1988; Begg and Doanachie, 1977; Mileykovskaya et al., 1998). Taken together, this suggests that switching from mid-cell to filamentation-based division may be a universal method for cells under stress to ensure viable cell divisions.

Consistent with previous reports (Wehrens et al., 2018), we find that damage-treated cells switch from mid-cell to polar division in a size-dependent manner. Our results suggest that the growth dynamics of filaments does not regulate size homeostasis at division. Instead, size of the daughter cell is determined by a combination of Min oscillations and chromosome segregation (Fig. S5). Cell growth may or may not occur to accommodate both these events, consistent with the observation that consecutive divisions can take place independent of addition of length in between, likely due to the accumulation of already duplicated termini that undergo segregation. Previous studies in wild type conditions have proposed that division timing could be modulated by factors such as rates of constriction or concurrency between processes of chromosome segregation and division (Micali et al., 2018; Lambert et al., 2018). In damage-induced filaments, we find that cell division is composed of two distinct steps of division site location and licensing. This results in a one-site-at-a-time cell division, which may serve as a checkpoint to ensure accurate and complete chromosome segregation. As seen for E. coli filaments under other stresses (Wehrens et al., 2018), the MinCDE system contributes to determining the location of divisome assembly. As cell length changes, the location of the Min oscillation nodes closest to the poles are likely to remain at a constant relative position, whereas the relative position of oscillation nodes away from the poles are likely to keep changing as the cell length changes. This would also explain why cells switch from mid-cell to poleward division as they increase in length. Subsequent to this, divisome assembly may be stabilized due to chromosome segregation away from mid-cell. Indeed, a recent study has shown that physical forces (such as molecular crowding) squeeze or confine DNA towards mid-cell in filaments (Wu et al., 2019), which could facilitate better or faster separation of chromosomes nearer the poles during segregation.

Our data are consistent with the idea that once the division machinery has assembled at a potential site of division, licensing occurs only when no DNA is encountered by the divisome; thus even in the absence of chromosome segregation anucleate cells can be produced (Mulder and Woldringh, 1989). However, upon initiation of chromosome resegregation, it is likely that a division checkpoint is triggered by the *terminus* region of the chromosome, via negatively regulating the ability of division to be completed. Consistently, we find that chromosome segregation can impose a delay in division, with the *terminus* being the last region of the chromosome to be segregated. Specifically affecting *terminus* segregation via deletion of *matP* results in perturbation to damage recovery, with a significant increase in anucleate cell production. These cells also have a broad distribution of nucleated daughter cell sizes. Hence concurrency between *terminus* segregation and cell division contributes to daughter size maintenance during asymmetric cell division. Given that MatP has two independent functions of *ter* region organization and *ter* anchoring with the divisome (Männik et al., 2016; Bailey et al., 2014; Nolivos et al., 2016), it would be important to understand which of these activities is specifically necessary in the recovery process. The system characterized in this study now provides the ability to assess these rate-limiting steps of division licensing and further probe the mechanisms of asymmetric chromosome segregation that preferentially results in daughter cells devoid of damage.

In sum, our study highlights the requirement for coordination between two independent mechanisms of divisome regulation for division accuracy in filaments. The Min system facilitates localization of FtsZ in nucleoid-free regions. In parallel, modulation of *terminus* region dynamics by MatP regulates the licensing of cell division. Concurrency between *terminus* positioning and segregation with cell division ensures daughter size maintenance in dividing filaments. Together, this results in the production of viable daughter cells with wild type growth and division dynamics and contributes to the restoration of cell size distribution, even when the filament itself may not recover. The conservation of asymmetric division in a range of stress-induced bacterial filaments underscores the importance of this mechanism in facilitating robust exit of cells from the DNA damage checkpoint via the generation of fit, damage-free daughter cells.

## Author contributions

SR: conception of project, experimental design, execution of experiments, data analysis and writing of manuscript. AC: experimental design and execution of experiments and analysis related to RecA. SK: data analysis and interpretation, writing of manuscript. AV: reagents, tools and interpretation. AB: conception of project, experimental design, writing of manuscript and funding.

## Acknowledgements

The authors are grateful to Thomas Bernhardt, Steven Sandler, Suckjoon Jun, Bill Soederstroem, Fred Boccard, Olivier Espeli, Christian Lesterlin, Ramanujam Srinivasan and Manjula Reddy for generous sharing of strains and plasmids. The authors acknowledge assistance from Ismath Sadhir in RNA extraction, Nitish Malhotra in RNA-seq analysis, Aditya Jalin, Alex Sam Thomas and Aalok Varma in writing Matlab scripts as well as the Central Imaging and Flow Facility (CIFF) and Next-generation genomics facility (NGGF). The authors thank Fagwei Si, Piet de Boer, Raj Ladher, Michael Laub, Tung Le and members of the AB lab for helpful discussions/ feedback on the manuscript. This work is supported by funding from NCBS-TIFR (AB and SK), by an HFSP Career Development award (AB) and funding from the Simons Foundation (SK).

## Materials and Methods

### Bacterial strains and growth conditions

Strains and plasmids used in the study are listed in Table 1. Transductions were conducted with P1 phage following the protocol in (Thomason et al., 2007). For *Escherichia coli*, cells were grown at 37°C in either rich media (LB: for 1 L dissolved 10 g tryptone, 5 g yeast extract and 10 g NaCl in doubled distilled water) or minimal media (M9-Cas: for 1 L dissolved 5 g glucose, 1 g casamino acids, 1 ml of 0.5% thiamine, 1 ml of 1M MgSO_4_ and 200 ml 5x M9 salts in double distilled water). DNA damage was induced with 1 μg/ml of mitomycin C (MMC) for 30, 60 or 90 min (LB) or 90 min (M9-Cas) (unless otherwise indicated). Cell division was inhibited using 5 μg/ml of cephalexin for 60 min (LB) or 90 min (M9-Cas).

### Fluorescence Microscopy

Imaging was performed on a widefield, epifluorescence microscope (Eclipse Ti-2E, Nikon with motorized xy-stage, Z-drift correction), 63X plan apochromat objective (NA1.41), pE4000 light source (CoolLED), OkoLab incubation chamber and Hamamatsu Orca Flash 4.0 camera. Images were acquired using the NIS-elements software (version 5.1). Microfluidics imaging was performed using the CellASIC-ONIX2 Microfluidic System, Temperature Controlled CellASIC-ONIX2 Manifold XT and CellASIC ONIX Plate for Bacteria Cells (B4A) (Merck). Details of the imaging and acquisition setting are described here (Raghunathan and Badrinarayanan, 2019).

### Time-course and Time-lapse imaging

For the recovery time-course, cells were grown overnight in LB or M9-Cas, back diluted to ∼OD_600_ 0.01 and allowed to grow to OD_600_ ∼0.1. Culture was then treated with MMC or cephalexin for 60 min (LB) or 90 min (M9-Cas). Cells were pelleted, washed with fresh media, resuspended to OD_600_ ∼0.1 and then allowed to recover from damage treatment. Cultures were maintained in log phase (OD_600_ ∼0.1 - 0.4) throughout the experiment. For time-course, samples for microscopy were collected at time points indicated. 1 ml of culture was pelleted, resuspended in 100 μl and then spotted on 1% agarose pad and imaged (Chimthanawala and Badrinarayanan, 2019).

For time-lapse experiments, damaged-induced cells were pelleted and washed with fresh media and were either loaded into the microfluidics device or spotted on a 1.5% agarose pad (made with appropriate growth media) and imaged. All time-lapse images were taken every 30 sec or 2 min for ∼3 hrs. For the FtsZ overexpression or FtsN imaging experiments, cells were induced with either 0.3% arabinose or 0.1% rhamnose respectively for 1 hr prior to imaging. For MinD-GFP imaging, cells were induced with 0.2mM IPTG for 90 min prior to imaging. Inducers were maintained in the agarose pads or microfluidic plates. Experiments were performed in LB unless otherwise indicated.

### RNA sequencing

Protocol for RNA extraction described in (Badrinarayanan et al., 2017) was followed. Briefly, cell pellets were collected during the recovery time-course at the indicated times. RNA was extracted using the Direct-zol™ RNA MiniPrep (Zymo, Cat no. R2052A81) and RNA Clean & Concentrator™-25 (Zymo, Cat no. R1018A82). RNA libraries were prepared using the TruSeq Stranded mRNA Library Preparation kit at NCBS NGS facility. Libraries were sequenced using Illumina MiSeq sequencing platform. Raw reads (single end; read length = 50 bp) were obtained as .fastq files. The reference genome sequence (.fna) and annotation (.gff) files for the same strain (accession number: NC_000913.3) were downloaded from the ncbi ftp website (“ftp.ncbi.nlm.nih.gov“). The raw read quality was checked using the FastQC software (version v0.11.5). BWA (version 0.7.12-r1039) (Burroughs and Aravind, 2016; Li and Durbin, 2009) was used to index the reference genome. Reads with raw read quality ≥20 were aligned using BWA aln -q option. SAMTOOLS (version 0.1.19-96b5f2294a) (Li et al., 2009) was used to filter out the multiply mapped reads. BEDTOOLS (version 2.25.0) (Quinlan and Hall, 2010) was used to calculate the reads count per gene using the annotation file (.bed) in a strand specific manner. Normalized counts per millions (cpm_n_) was obtained using TMM normalization of EdgeR package. The cpm_n_ across time was calculated relative to the no damage control. The log_2_ fold change in expression was plotted from this.

### Image Analysis

Images acquired were visualized and processed using ImageJ (Schindelin et al., 2012, 2015). Segmentation was performed using Oufti (Paintdakhi et al., 2016). Foci tracking was accomplished using the Spotfinder Z function of MicrobeTracker (Sliusarenko et al., 2011). For recovery time-course, custom Matlab scripts were used to extract cell length and total fluorescence intensity information for each cell. Anucleate cells were identified using a threshold of 0.02 a. u. for total intensity of HupA-GFP in the cells. For time-lapse analysis, at each division, the longer cell was termed L_D_ and the smaller cell given the identity of S_D_. A custom Matlab script was used to the extract the lengths of the L_D_ and S_D_ at each division, the length added between divisions and time between divisions. Daughter cells were classified as ‘recovers’ if at their first division, their length was <10µm and they divided symmetrically. Cells that were longer at their first division and divided asymmetrically were classified as ‘filaments’. RecA foci numbers were obtained using Spotfinder Z and combined with cell length information from Oufti. Number of FtsZ rings in a cell were obtained using Spotfinder Z and combined with cell length information from Oufti. For analysis of constrictions and divisions, we used Oufti, which allows sub-pixel segmentation of phase contrast images of cells in a time-lapse. The method is illustrated in figure S2 in the supplementary methods section. In (a), phase profile for a wild type cell before and after division is plotted. While the profile is, on average, a flat line for most of the time imaged, one can identify a constriction prior to division (marked with *). Similarly in (b), the phase profile for a filamentous cell (DNA damage-induced) undergoing division is plotted. If the phase profile deviates by 20-25% of the cell width, it is called a constriction. If the phase profile is discontinuous (with a gap), then it is marked as a division by the segmentation algorithm automatically. For nucleoid tracking, fluorescence intensity profiles of HupA-mCherry, obtained from Oufti was used. Data was first smoothened using a Savitzgy-Golay filter (Luo et al., 2005). Following this the fluorescence profile was inverted so that the regions of lowest fluorescence now had the highest values. Peaks were determined using the peak prominence function after setting a threshold of half the maximum intensity for each frame. Peaks correspond to lowest fluorescence intensity in the cell. Regions right next to the cell poles were excluded. This information was combined with FtsZ foci tracked using Spotfinder Z to obtain relative positions of the lowest fluorescence intensity in the cell and FtsZ ring. Formation of FtsZ, ZapA and FtsN foci, and segregation of the single *terminus* focus (ParB-GFP) to 2 foci at the site of division was scored manually along with time to division.

